# Stable topical application of antimicrobials using plumbing rings in an *ex vivo* porcine corneal infection model

**DOI:** 10.1101/2024.12.02.626328

**Authors:** Daniel M. Foulkes, Keri Mclean, Taruni Masharma, David G. Fernig, Stephen B. Kaye

## Abstract

Microbial keratitis (MK) is a substantial cause of clinical blindness worldwide. *Pseudomonas aeruginosa* is an opportunistic gram-negative bacterium and is the leading cause of MK. Infection models are vital tools in understanding host-pathogen interactions and the development of novel therapies. As well as ethical and practical advantages, *ex vivo* infection models can allow researchers to better investigate host-pathogen interactions more accurately than traditional cell culture systems. The versatility of porcine corneal *ex vivo* models have been employed to study various pathogens (for example *Staphylococcus aureus* and *Acanthamoeba*) and has enabled innovation of novel MK therapies. Here, we describe an improved porcine corneal *ex vivo* protocol, which uses plumbing rings and medical adhesive to circumvent several distinct limitations and challenges. The application of a 10 mm plumbing ring to the center of the cornea allows localized inoculation of pathogens of interest, maintaining them at the site of infection, rather than running the risk of “run off” of topically added aqueous solutions. The second important advantage is that topically applied therapeutic agents can be properly maintained on the cornea within the plumbing ring reservoir, allowing more accurate study of antimicrobial effects. In this contextualized protocol, we infected porcine corneas with the *P. aeruginosa* strain PA103 with topical treatments of moxifloxacin. PA103 colony forming unit (CFU) deduction, quantification of corneal opacity and histology analysis (hematoxylin and eosin staining), was used to assess infection over 48h, which was alleviated by moxifloxacin in a dose-dependent manner.

## 1. Introduction

Corneal opacification is the sixth leading cause of global blindness, accounting for 3.2% (0.5%-7.2%) of 36 million cases. Microbial keratitis (MK) is the main cause of corneal opacification. The corneal epithelium maintains a barrier to invading microorganisms, however, infection may occur when this barrier is disrupted by trauma, contact lens wear or surgery [1]. MK normally presents through a painful epithelial defect with associated signs of corneal stromal inflammation and ulceration. There is often a localized corneal opacity with epithelial erosion, accompanied by an anterior uveitis, fibrinous exudate or a hypopyon. Infection may progress to a descemetocele and perforation and occasionally MK can cause whole eye loss, which is more frequently observed in older patients [1].

*Pseudomonas aeruginosa* is an opportunistic gram-negative pathogenic bacterium and accounts for 25% of MK cases. *P. aeruginosa* possesses an array of virulence factors, although only some are implicated in MK. The type III secretion system (T3SS) is a fundamental driver of *P. aeruginosa* pathogenicity, of which the two main effectors are Exotoxin U (ExoU) [2, 3] and Exotoxin S (ExoS) [4, 5]. ExoU expression is associated with increased MK disease severity with poor clinical outcomes. The ExoU phospholipase is injected into host cells via the T3SS, causing rapid destruction of host cell plasma membranes [2]. ExoS modulates the cytoskeleton of host cells by

ADP-ribosylating specific host targets such as Ras and Rho proteins, facilitating bacterial invasion and intracellular survival of *P. aeruginosa* in chronic infections [6]. Several key secreted proteases of *P. aeruginosa* also serve important roles during infection. Elastases (LasA and LasB), protease IV, alkaline protease and *P. aeruginosa* small protease (PASP) have distinct host targets that contribute to tissue damage, degradation of extracellular matrix (ECM) components [7-9] and subversion of the host immune system through targeting of neutrophils, complement proteins, immunoglobulins, antimicrobial peptides, host extracellular enzymes and surfactants [8, 10-14].

Commissioned by the UK parliament, it is predicted in O’Neill’s Review on Antimicrobial Resistance that by 2050, deaths resulting from infections by antibiotic resistant bacteria will surpass deaths from cancer [15]. The alarming efficiency in which bacteria have developed resistance against our existing therapeutic arsenal of antimicrobial agents, means that there is an urgent need to develop new therapies. Due to their accessibility and with ethical considerations (reducing dependency on extensive *in vivo* studies), *ex vivo* models are becoming increasing popular tools to interrogate mechanisms of infection and assess novel therapeutics. Porcine eyes are inexpensive (our lab is charged £2 per eye) and are available from local abattoirs. From histological and immunohistochemical characterization, it has been demonstrated that, despite the apparent absence of a Bowman’s layer, the porcine corneal structure and molecular markers closely resembles that of human’s, despite possessing greater thickness (1100-1497 µM for porcine and 500-550 µM for human) [16, 17]. This increase in thickness, however, means that porcine corneas possess a greater available area for pathophysiological investigations. Beyond MK, the *ex vivo* cornea is a valuable testbed to assess infection mechanisms and test novel therapeutics, as it is an accessible, robust and transparent organ, which may also give researchers the opportunity to translate their findings to other infections sites.

In recent years, there have been various developments and optimizations in the use of *ex vivo* porcine eye models for the testing of new therapies. Kennedy *et al*. (2020) developed an *ex vivo* porcine corneal *P. aeruginosa* infection model to test the efficacy of patented poly-epsilon-lysine peptide hydrogels in the treatment of MK [18]. Whole porcine corneas were dissected with surgical scissors and scalpel, placed epithelial side down into sterile bijou tube lids with agarose dissolved in Dulbecco’s modified eagles’ medium (DMEM) then added to produce a cornea with agarose support. When placed epithelial side up in a 6-well plate, pathogen and/or therapeutic agent can then be added to the corneal dome. It is important to note that to achieve consistent infection, the epithelial barrier must be first damaged. Some studies opt for the debriding of epithelial cells by application of 70% ethanol [18, 19], while others report mechanical trauma with a scalpel or needle tip (30G) to be effective [20, 21]. With both bacteria and therapeutic agent being topically added to the cornea, there is significant risk that aqueous solutions run from the center of the cornea into the 6-well plate, precluding experimentation. To circumvent this challenge, Okurowska *et al* (2020) developed a protocol that employs custom made glass molds that fit around the whole cornea with a 10 mm opening at the top in which to add aqueous solutions [20]. Although these bespoke glass molds are innovative, their production requires a skilled glass blower within a specialized facility.

In the present study, we adapted the protocol of Kennedy *et al* (2020) to include the attachment of plumbing rings (readily available from hardware stores), with medical adhesive, to the surface of the cornea. The plumbing rings act as a reservoir whereby an aqueous volume of up to 150 µL can be added and maintained in a centrally localized position on dissected porcine corneas. As proof of concept, we infected *ex vivo* porcine corneas with the cytotoxic, ExoU producing, strain of *P. aeruginosa*, PA103, and observed effects of infection after 48h. Moxifloxacin is a fluroquinolone antibiotic that is often used to treat *P. aeruginosa* MK. Therefore, we topically applied varying concentrations of moxifloxacin to infected porcine corneas to test the efficacy of an established MK therapeutic in the developed *ex vivo* model. Measurements included, imaging of corneas, quantification of opacity, deduction of PA103 colony forming units (CFU) in homogenized corneas, and histological analysis of sectioned corneas by hematoxylin and eosin (H&E) staining.

## 2. Materials and methods

### 2.1 Materials

The following materials and reagents were required for this study: whole pig eyes obtained the morning of slaughter (CS Morphet’s abattoir, Widnes, UK), surgical scissors, forceps and scalpel (size 10, Fisher scientific), phosphate-buffered saline (PBS), 10% (v/v) iodinated povidone (Ecolab), ethanol, 10 mm Plumbsure Rubber O rings (B&Q, Bidston, UK), LIQUIBAND () 0.8 g Exceed topical skin adhesive, Luria– Bertani (LB) broth, LB agar (Melford), UltraPure Agarose (Thermo Fisher), DMEM with 10% (v/v) fetal bovine serum (Labtech), 12-well plates, moxifloxacin (Merck) and H&E staining kit (Abcam).

### 2.2 Preparation of bacterial inoculum

The *P. aeruginosa* strain PA103 was cultured in 100 mL of LB broth, from a glycerol stock, overnight at 37ºC in a shaker incubator. 2 mL of the PA103 overnight culture was then added to 20 mL of fresh LB broth and expanded to an OD_600_ of 0.8. Of this subculture, 1 mL was taken and centrifuged for 5 minutes at 5000 **g**. The supernatant was discarded, and the bacterial pellet resuspended in 10 mL of PBS, yielding a PA103 suspension at a concentration of 10^7^ CFU/mL.

### 2.3 Preparation of porcine corneas

The eyes of large white landrace pigs were removed the morning of slaughter and transferred to PBS. In a laminar flow hood, using forceps to hold the eyes in place, a small incision was made at the edge of the sclera, releasing ocular pressure. Surgical scissors were then used to cut around the cornea, whilst maintaining ∼2.5 mm of sclera outside the circumference of the cornea in a full circle. The resulting corneoscleral button was separated from the uveal tissue using two sets of forceps. In a 10 cm dish, corneas were subsequently incubated in 10% iodinated povidone for 2 minutes for sterilisation. Corneas were then washed 3 times in PBS. In a sterile 10 cm dish, one side of 10 cm plumbing rings were immersed onto LIQUIBAND topical skin adhesive. Forceps were then used to compress all edges of the ring to the center of the cornea, ensuring a tight seal.

Corneas were placed epithelial side down. UltraPure Agarose (0.5% [w/v]) was dissolved in DMEM in a 65.5 °C water bath, cooled to approximately 37 °C and then pipetted onto the endothelial side of corneas to fill the cavity. Corneas with solidified agarose supports were transferred into 12-well plates, epithelial side up.

To induce a controlled epithelial wound on the cornea, 10 µL 70 % ethanol (v/v in PBS) was applied to the center of the corneal surface for 10 seconds. Following ethanol exposure, the epithelial layer was debrided using the edge of a scalpel around the treated area to simulate mechanical injury. In the 12-well plate, the corneas were rinsed three times with PBS to ensure complete removal of ethanol and debris. Finally, 1 mL of DMEM was added to each well, covering the outer edge of the corneal epithelium without submerging the plumbing ring.

### 2.4 Inoculation of corneas and topical application of MK treatment

For each cornea that was infected, 10 µL of a 10^7^ CFU/mL suspension of PA103 in PBS was added to Eppendorf tubes with the concentrations of moxifloxacin indicated in the figure legends. The final volume of each suspension was made up to 150 µL with PBS, which was then added to the plumbing ring reservoir at the centre of the cornea. This equated to an addition of 1 x10^5^ CFU of PA103 inoculated onto each cornea. Corneas, in12-well plates, were transferred to a 37 °C incubator in 5 % CO_2_ (v/v) for 48 h.

### 2.5 Determination of CFU within corneas

After 48 h infection, corneas were gently separated from the agarose supports and plumbing rings and then washed in PBS. Corneas were then transferred into 500 μL PBS and homogenized using a Qiagen Tissue ruptor (Qiagen). Suspended bacteria were serially diluted in PBS (10^−1^-10^−6^) with 10 µL plated onto LB agar plates followed by overnight incubation at 37°C. After incubation, colonies were counted on plates with bacterial load calculated by multiplying the counted colonies by dilution factor and the plated volume.

### 2.6 Imaging and spectrophotometric analysis of corneal opacity

After 48 hours of infection, corneas were removed from agarose supports and plumbing rings and then rinsed in PBS. To assess corneal opacity visually, corneas were imaged using a BioRad ChemiDoc system, capturing optical density data. For quantitative analysis, corneas were transferred to a 12-well plate, and opacity was measured using a Hidex spectrophotometer by evaluating transmitted light permeability at 400 nm.

### 2.7 Histological analysis of porcine corneas

Corneas were processed using a Leica ASP300 tissue processor. Tissues were fixed in formalin, dehydrated through a graded series of ethanol, cleared in xylene, and then embedded in paraffin wax. Five μm sections of paraffin-embedded corneal tissues were cut using a microtome and mounted onto glass slides.

For histological analysis, the tissue sections were stained with H&E using a staining kit (Abcam), following the manufacturer’s protocol. Once stained, the sections were imaged using a VENTANA DP 200 slide scanner (Roche). The open-source software QuPath was used to view and analyse images for morphological evaluation of corneal tissue and to assess the extent of infection and tissue damage.

## 3. Results and discussion

We tested the efficacy of moxifloxacin to reduce PA103 infection of porcine corneas, *ex vivo*. In our developed protocol, PA103 inoculum and aqueous moxifloxacin was maintained on the cornea with use of a plumbing ring reservoir, employing the preparatory method described in figure 1. Downstream analysis consisted of corneal CFU count, corneal opacity and histology analysis.

**Figure 1:**
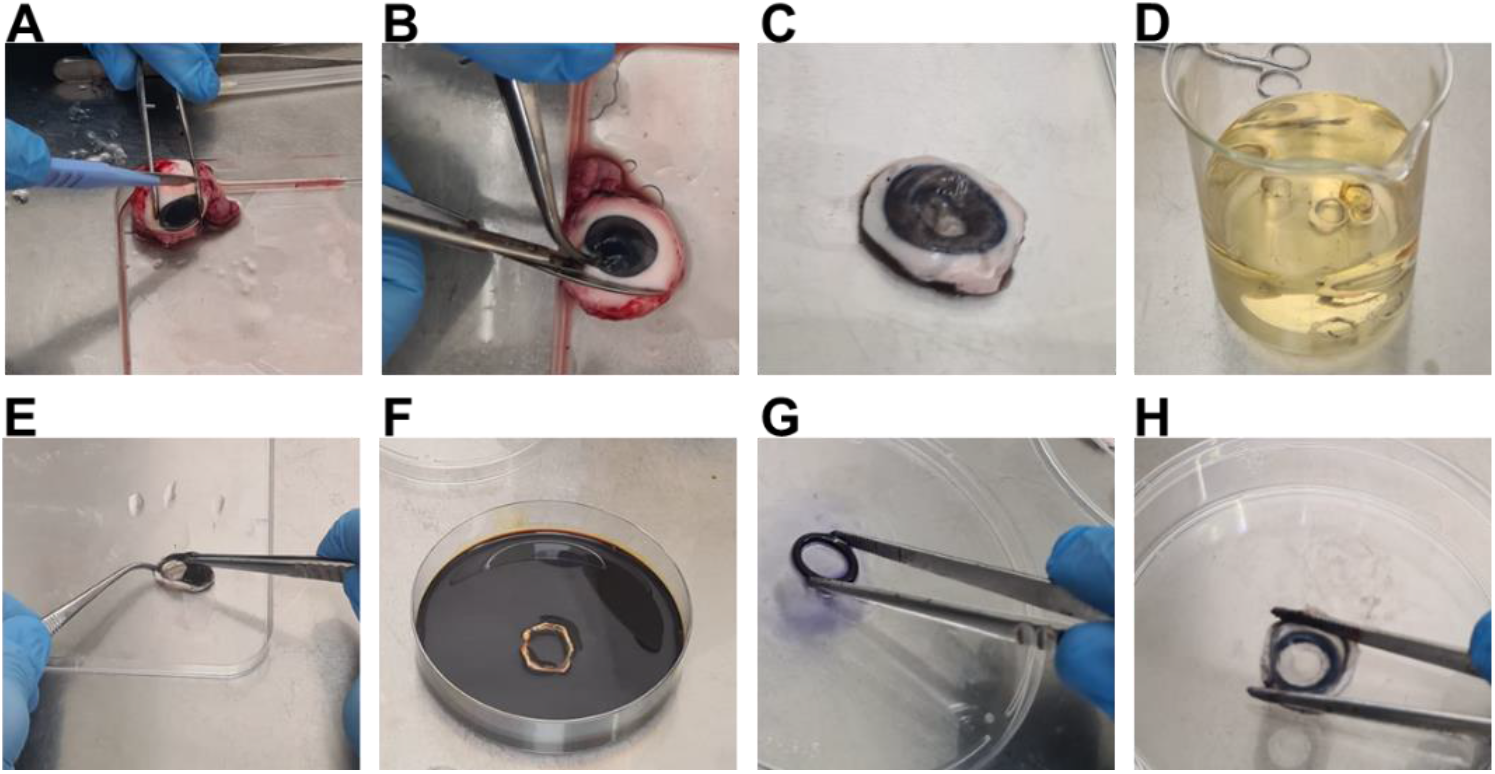
Application of plumbing ring during porcine cornea preparation. (A) A scalpel was used to make an incision into the sclera. (B) Scissors were used to cut around the cornea to yield (C) a corneal scleral button. (D) Two pairs of forceps were used to remove the uveal tissue. (E) The corneas were immersed in 10 % (v/v) iodinated povidone for 2 minutes and then washed 3 times in PBS (F). (G) LIQUIBAND adhesive was applied to a sterile 10 cm petri dish, and a 10 mm plumbing ring was immersed in this. (H) Using forceps, the epithelial side of the porcine corneas was pressed onto the adhesive-coated side of the plumbing ring, ensuring an even seal.

### 3.1 CFU analysis to determine efficacy of moxifloxacin in PA103 infected Porcine Corneas

PA103-infected corneas, in the absence of moxifloxacin, exhibited a bacterial expansion from an initial inoculum of 5 log_10_ CFU to 10.5 ± 0.6 log_10_ CFU after 48 h of infection (Figure 2). This level of *P. aeruginosa* growth aligns with findings from previous studies [20]. Prior research has demonstrated that *P. aeruginosa* reaches a maximum growth of approximately 10^9^-10^10^ CFU within 24 hours post-infection, with extended incubation (48 hours) showing no significant further increase in bacterial load. This plateau in growth after prolonged incubation is likely due to nutrient depletion, which limits further bacterial expansion.

**Figure 2:**
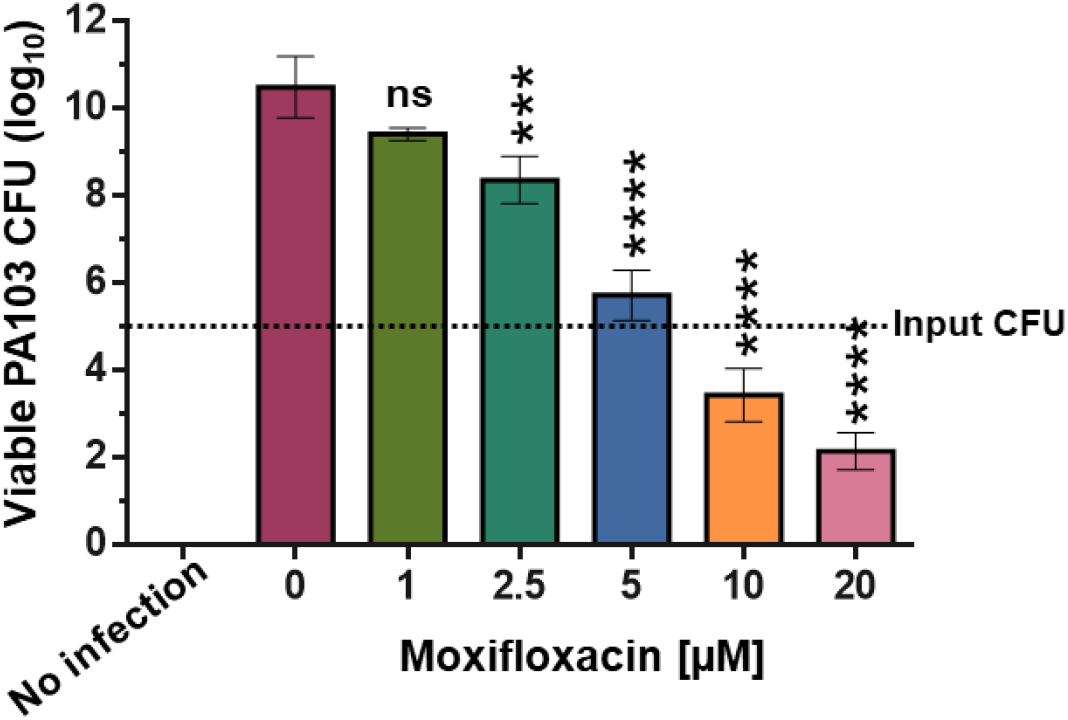
PA103 CFU detection in *ex vivo* porcine corneas after 48h infection and moxifloxacin treatment. *Ex vivo* porcine corneas were infected with 1 × 10^5^ CFU PA103 for 48 h in the indicated concentrations of moxifloxacin. Whole corneas were homogenized, diluted in PBS followed by plating, overnight incubation at 37 ºC prior to CFU counting. Results were plotted and one-way ANOVA with multiple comparisons (to 0 µM moxifloxacin) performed in Prism ((*<0.05, **<0.01, ***<0.001, ****<0.0001).

By administering increasing concentrations of moxifloxacin to PA103-infected corneas, we observed a dose-dependent reduction in viable CFU recovered from homogenized corneas (Figure 2). Our previous *in vitro* studies determined the minimal inhibitory concentration (MIC) of moxifloxacin for PA103 growth to be 4 µM [22]. Although significantly lower than non-treated controls, at 5 µM moxifloxacin, there was a slight increase in detectable PA103 CFU compared to the initial inoculum. Even at 5 times the *in vitro* MIC (20 µM), while a significant reduction was observed, 2.1 ± 0.3 log_10_ CFU of viable PA103 persisted. This is consistent with recent findings where the related fluoroquinolone, ciprofloxacin, failed to completely eliminate *P. aeruginosa* from *ex vivo* porcine corneas, even at concentrations above the MIC [23]. The concentrations of moxifloxacin used in this study are substantially lower than those typically employed in clinical settings, potentially explaining the incomplete clearance of PA103 from the corneas. Additionally, factors such as antibiotic bioavailability and penetration in *ex vivo* porcine corneas, as well as the absence of fully operational host immune defenses, may also contribute to the observed persistence of bacteria.

### 3.2 Measurement of corneal opacity to determine moxifloxacin effectiveness

Topical solutions of moxifloxacin added to PA103-infected *ex vivo* porcine corneas resulted in a dose-dependent reduction in corneal opacity after 48 hours, as measured by optical density at 400 nm (Figure 3). In the absence of moxifloxacin, a significant increase in opacity was observed, consistent with severe infection and corneal damage caused by *P. aeruginosa*. Treatment with moxifloxacin at 1, 2.5, and 5 µM led to modest decreases in opacity, suggesting limited efficacy in mitigating infection-related corneal clouding at these concentrations. However, 10 µM moxifloxacin significantly reduced opacity, while 20 µM restored corneal transparency to levels comparable to uninfected controls.

**Figure 3:**
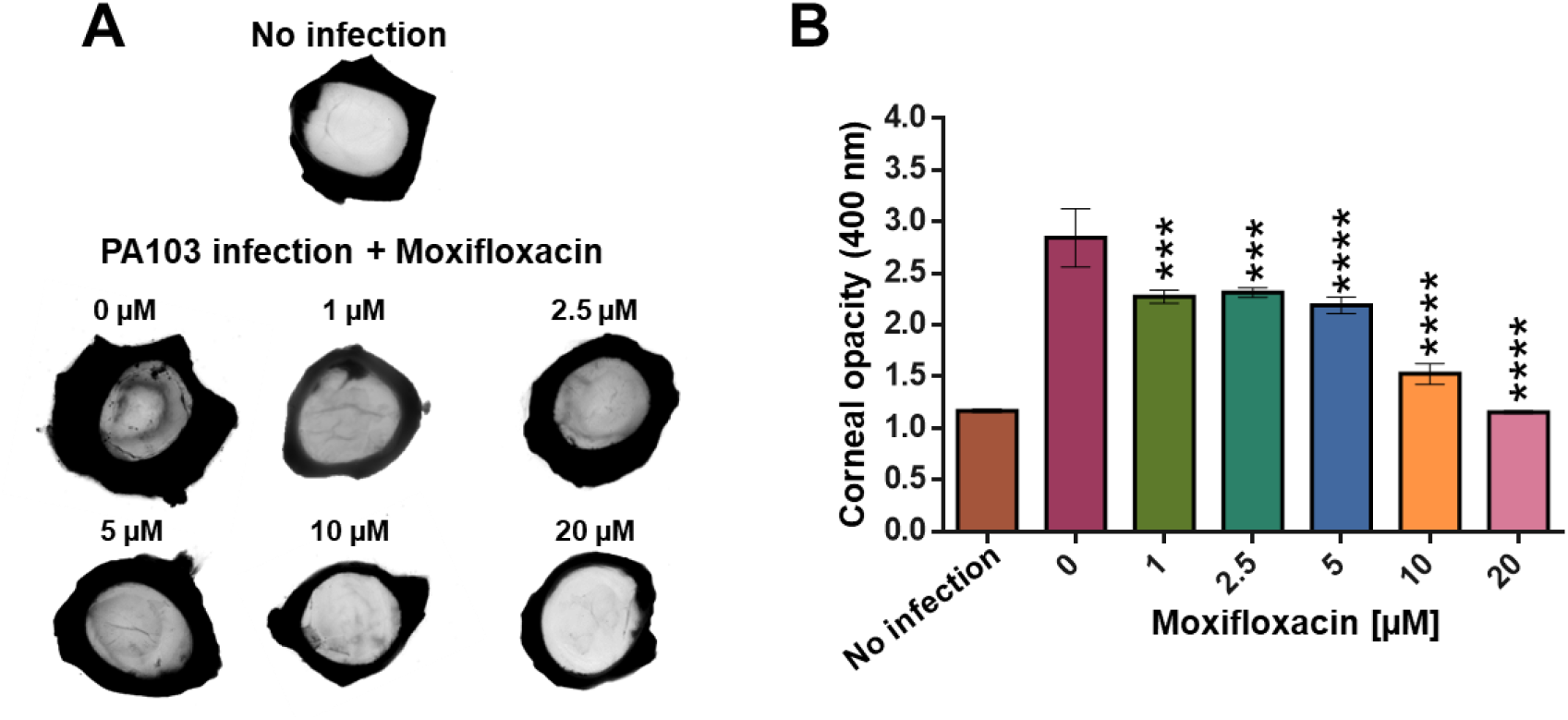
Opacity measurement to determine moxifloxacin efficacy in PA103 infected *ex vivo* porcine corneas. *Ex vivo* porcine corneas were infected with 1 × 10^5^ CFU PA103 for 48 h prior to (A) scanning for image acquisition and (B) optical absorbance measurement at 400 nm in a 12-well plate using a spectrophotometer. Results were plotted and one-way ANOVA with multiple comparisons (to 0 µM moxifloxacin) performed in Prism (*<0.05, **<0.01, ***<0.001, ****<0.0001).

Despite the restoration of corneal clarity at 20 µM, viable *P. aeruginosa* were still detected, as evidenced by CFU counts (Figure 2). The absence of corneal opacity at this concentration, despite the persistence of bacteria, may be explained by the antimicrobial action of moxifloxacin in effectively suppressing bacterial virulence and toxin production, thereby preventing further tissue damage and clouding. Previously, we demonstrated, *in vitro*, that moxifloxacin concentrations greater than 4 µM reduced PcrV expression [22], a critical component of the T3SS. Additionally, moxifloxacin may have reduced the bacterial load to a level that, while not entirely eradicating the infection, was insufficient to induce significant inflammatory responses or tissue degradation associated with corneal opacity. This suggests that while moxifloxacin at 20 µM was able to preserve corneal integrity, complete bacterial clearance may require higher concentrations or additional treatments to fully eliminate the pathogen.

### 3.3 Histology to evaluate PA103 infection after moxifloxacin treatment

Using histology, with H&E staining, it is possible to assess the structural changes and erosion that occurs to the cornea during *P. aeruginosa* infection. Typically, during infection, cellular damage, thinning and ulceration (erosion) can be observed in the epithelium (Figure 4, black arrows). In the stroma, edema can be observed with increased space between collagen fibrils (Figure 4, white arrows), which can be quantified, if necessary. It is also possible to employ gram staining to observe quantity and penetration of infection [19].

**Figure 4:**
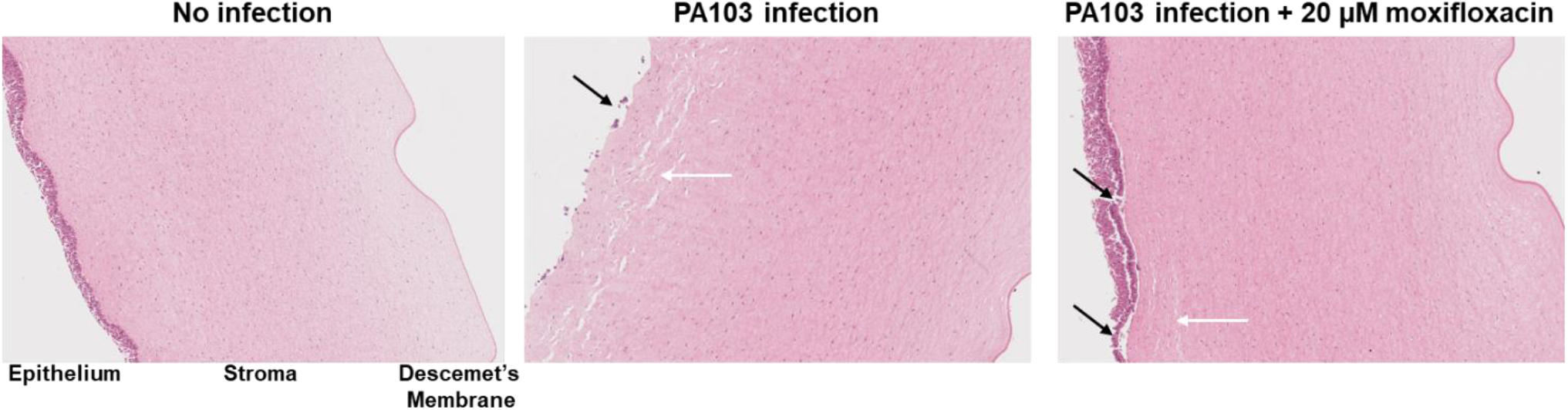
Histological analysis of PA103 infected porcine *ex vivo* corneas. *Ex vivo* porcine corneas were infected with 1 × 10^5^ CFU of PA103 for 48h with and without 20 moxifloxacin. After parafilm embedding and sectioning, slides were stained with H&E. Black arrows indicate epithelial ulceration and white arrows stromal edema.

Without infection, regular and undamaged epithelial layers with intact stroma could be observed (Figure 4). After 48h PA103 infection, we observed complete epithelial erosion and extensive stromal edema (Figure 4). With 20 µM moxifloxacin, the corneal epithelium was mostly undisrupted with only partial stromal edema. In all instances, the Descemet’s membrane remained undamaged, which would otherwise indicate extensive infection and complete corneal breakdown. We note that it is not usually possible, due to processing limitations, to retain the squamous endothelium, which resides beyond the Descemet’s membrane.

## 4. Limitations

### 4.1 Porcine cornea physiology considerations

While porcine corneas share many anatomical and physiological similarities with human [16, 17], certain differences, such as the greater thickness of porcine corneas, may influence the progression of infections and the permeability of therapeutic agents. This difference could lead to altered permeation rates and retention times for drugs in porcine corneas, particularly for hydrophilic drugs, which face more resistance crossing the denser stroma [24]. This discrepancy could impact the direct translation of findings to human ocular infections. Moreover, unlike human corneas, porcine corneas lack a Bowman’s membrane, a structure that serves as a protective barrier and contributes to resistance against injury and infection. However, porcine corneas have evolved to maintain transparency and structural integrity, potentially compensating for the absence of the Bowman’s membrane through additional epithelial cell layers. Although the *ex vivo* porcine model offers a valuable platform for studying infection mechanisms and testing novel treatments, it is crucial to validate findings in *in vivo* models. This step is essential to assess therapeutic efficacy in a more complex biological environment before considering clinical applications.

### 4.2 Limitations of the *ex vivo* environment

Despite its advantages, the porcine eye *ex vivo* model has distinct limitations. The lack of a fully functional immune system in this model excludes innate and adaptive host immune responses, which play a critical role in infection control and healing *in vivo*. Thus, although resident macrophages and low numbers of neutrophils are present in the cornea prior to infection [25], the lack of a circulatory system in *ex vivo* models limit its ability to fully replicate the complex interactions between bacteria and the host’s immune response. This limitation likely contributes to the incomplete eradication of PA103 by moxifloxacin after 48 hours (Figure 2). Additionally, the restricted availability of nutrients in *ex vivo* systems may alter bacterial behavior and treatment efficacy, further complicating the model’s ability to fully mimic *in vivo* infections.

The absence of tear film, which contains antimicrobial peptides, enzymes, immunoglobulins and surfactants, removes an essential component of the eye’s natural defense. Further developments to the porcine *ex vivo* cornea model could consist of the incorporation of artificial tear solutions, containing components such as electrolytes (NaCl, KCl, MgCl_2_), lysozyme, lipids, and mucins [26, 27]. *In vitro* these are used to mimic natural tear fluid and are often used in drug permeability studies. Advanced *in vitro* models, such as tear film lipid layer and microfluidic tear film models might be incorporated into *ex vivo* experimental design to study drug dissolution, and pathogen interaction with the ocular surface.

### 4.3 Experimental design considerations

Certain experimental limitations must be carefully considered during experimental design. Previous studies have shown that even a low initial inoculum of *Pseudomonas aeruginosa* (cytotoxic strain PA14), as few as 215 CFU, is sufficient to establish infection in *ex vivo* porcine corneas [23]. Remarkably, the bacterial load consistently reached a maximum of approximately 1 × 10^9^ CFU after 18 hours of infection, regardless of the initial inoculum size (ranging from 215 to 1 × 10^7^ CFU). While inoculating with such a low CFU count can enhance the dynamic range for observing bacterial load reductions in response to treatments, it also introduces potential variability due to the challenges of accurately mixing and pipetting at these low concentrations. Therefore, careful consideration of the initial inoculum size is essential, balancing the need for sensitivity in detecting treatment effects with the desire to minimize experimental variation.

In our experience, endothelial cells are not consistently recovered for histological analysis following dissection. The corneal endothelium is a single layer of cells that is crucial for maintaining corneal hydration by regulating fluid exchange, but these cells are delicate and easily damaged during handling and processing, which might present limitations when studying highly penetrant infections. While the *ex vivo* porcine eye model allows for the evaluation of the integrity of the Descemet’s membrane, the inability to reliably analyze the endothelium hinders comprehensive investigation of deeper corneal infections. In cases involving corneal perforation or complete corneal melting, histological analysis is further compromised, preventing detailed insight into the fate of the endothelial layer. As a result, this model may not fully capture the extent of endothelial damage in severe or advanced infections, necessitating complementary approaches to fully understand these processes.

## 5. Conclusions

In this study, we describe our advances in using the porcine eye *ex vivo* model as a simple, robust, reproducible, and cost-effective method for investigating ocular infections. A key challenge we addressed was the tendency for bacterial inoculum and topically applied treatments to run off the corneal epithelium, preventing consistent infection and treatment delivery. By incorporating plumbing rings, we were able to localize both the bacterial inoculum and therapeutic compounds, covering a substantial portion of the corneal surface while maintaining consistent exposure. Prior to this modification, it was difficult to reliably achieve reproducible *P. aeruginosa* infections in porcine corneas when applying aqueous therapeutics. While the isolated application of small, stable volumes (5-10 µL) directly onto the center of the cornea is sufficient for testing solid agents like hydrogels or contact lenses [18, 19], larger aqueous volumes often could not be retained on the corneal surface. This led to uncertainty regarding the effective concentration of the applied treatment and how much compound remained in contact with the cornea. Using moxifloxacin as a standard treatment, in this contextualized protocol we demonstrate the effectiveness of moxifloxacin to reduce corneal opacity, epithelial ulceration and stromal edema, highlighting the use of the developed *ex vivo* porcine eye model to assess topical therapeutics for bacterial infection.

